# A temperature-dependent shift in dietary preference alters the viable temperature range of *Drosophila*

**DOI:** 10.1101/059923

**Authors:** M Brankatschk, T Gutmann, M Grzybek, B Brankatschk, U Coskun, S Eaton

## Abstract

How cold-blooded animals adapt their behaviour and physiology to survive seasonal changes in temperature is not completely understood - even for well-studied model organisms like *Drosophila melanogaster*. Here, we show that *Drosophila* can extend their viable temperature range through temperature-dependent changes in feeding behaviour. Above 15°C, *Drosophila* feed and lay eggs on yeast. In contrast, below 15°C, *Drosophila* prefer to feed and lay eggs on plant material. The different lipids present in yeast and plants improve survival at high and low temperatures, respectively. Yeast lipids promote high tempera-ture survival by increasing systemic insulin signalling. This expands the range over which developmental rate increases with temperature, suggesting that faster nutrient utilization is required to fuel biochemical reactions driven faster by ki-netic energy. In addition to speeding development, yeast lipids increase fertility. Thus, yeast provide cues that could help Drosophila to exploit a transient summer food resource. Plant lipids, on the other hand, are required to maintain mem-brane lipid fluidity at low temperature, and increase cold-resistance of larvae and adults. The cold-resistance and lowered insulin signalling conferred by feeding on plants allows adults to survive for many months at temperatures consistent with overwintering in temperate climates. Thus, temperature-dependent changes in feeding behaviour produce physiological changes that could promote seasonal adaption.

---

Cold-blooded animals (poikilotherms) can undergo a wide range of fascinating metabolic, physiological and developmental changes that help them cope with seasonal and diurnal temperature fluctuations. Throughout some of the viable temperature range, these animals are able to complete an entire life cycle, raising the fascinating question how cellular and organismal processes underlying growth and development are coordinated at different temperatures. At temperature extremes, poikilotherms can enter dormant states that are resistant to temperature-induced damage. Responses to cold temperatures include the production of anti-freeze proteins, increases in polyols such as sugars and proline that reduce the freezing point(*1, 2*), and changes in lipid composition that maintain membrane fluidity - a phenomenon known as homeoviscous adaption(*3*). The efficiency of this compensation in fishes has been reported to vary between 20% and 100% depending on species and tissue type, and is associated with changes in fatty acid saturation and the PE/PC ratio. Goldfish show a > 20% difference in fatty acid unsaturation, and the mole% PC decreases 50% (4–6).

On the other end of the spectrum, high temperatures cause different problems. In poikilotherms, the metabolic rate increases with temperature, so they need more energy to develop and maintain a given body size(*7, 8*). This may account for the observation that animals that develop at higher temperatures mature at a smaller body size(*9*). Increasing temperature also speeds the rate of development - at least within the range over which development is successful(IO). Whether poikilotherms actively increase their rate of development at high temperatures is unknown - one explanation for faster development at high temperature is simply that chemical reactions are driven faster, as described by the Arrhenius equation(*10–13*). Even rates of biological “reactions” like cell division(*14*) or insect locomotion(*15*) respond to temperature in this way over a large range. But it is clear that the capacity to increase the organismal developmental rate with temperature is limited. The temperature-dependence of organismal development deviates significantly from an Arrhenius relationship(*13*). When the temperature becomes too high, the rate of development begins to slow down and then fail. It is not clear which biological processes set this limit.

*Drosophila melanogaster* provides a powerful genetic system in which to probe the mechanisms underlying temperature adaption. There has been extensive work on the ecology of this organism, and the genetic regulatory networks underlying its growth and metabolism have been well studied in the lab. Increasing the synergy between ecological and molecular genetic approaches could deepen our understanding of *Drosophila* ecology, and also provide a more realistic context in which to interpret laboratory studies, which often rely on temperature shifts to regulate gene expression.

Populations of *Drosophila melanogaster* are thought to survive year-round in temperate and tropical climates and are found in habitats with wide range of average seasonal temperature fluctuations(*16*). The response of *Drosophila* adults to temperature extremes has been studied using *Drosophila* stocks recently established from wild populations and then maintained on standard lab diets. While successful development is restricted to the range between 12°C and 30°C, adults can withstand a few hours at higher (38°C) and lower (−2°C) temperature extremes(*16, 17*). These limits can vary slightly depending on the geographical area from which the flies are isolated, and can be somewhat extended by “hardening”, i.e. pre-exposure to high or low temperatures(18–20). Since temperatures even in temperate climates can exceed these limits, it seems likely that *Drosophila* must behave in a way that enables them to avoid extreme temperatures in their natural environments, or that key ingredients in heat or cold adaption are missing from the lab, or both.

The physiological changes that occur in *Drosophila melanogaster* during rapid cold hardening have been explored using metabolomics, proteomics and transcriptom-ics(*21–25*). There is no evidence that *Drosophila* produce antifreeze proteins. Several studies report increases in glucose and trehalose levels in response to cold(*21, 23*), although this is not always seen(*18, 26*). The extent to which homeoviscous adaption occurs in *Drosophila melanogaster* has been controversial. Fluidity at different temperatures has not been directly measured in *melanogaster*. Small changes in fatty acid unsaturation have been noted upon rapid cold hardening or other forms of temperature ac-climation(*27–30*), however the differences that are observed are smaller than in other cold-blooded animals(4–6). Some(*29–32*) but not all(*26, 33*) studies report temperature-dependent changes in the ratio of PE to PC.

At high temperatures, *Drosophila* increase their metabolic rate like other poikilo-therms(*34*). Similarly, high temperatures speed development and produce smaller ani-mals(*13, 35*). The hormonal networks that control growth and developmental progression have been extensively studied, but how they interface with temperature is not clear.

Recent work in our lab led us to wonder whether diet might influence how *Drosophila melanogaster* respond to temperature changes. In the wild, these animals are thought to feed primarily on the yeasts present on decomposing fruit. Olfactory and gustatory cues attract *Drosophila melanogaster* to yeasts, both as a food source and as a substrate for egg laying(*36–41*). It is unclear to what extent plant material normally contributes to their nutrition. We recently developed two different food recipes containing exclusively yeast or plant material. Although these diets differ in specific ingredients, they contain similar proportions of calories derived from protein, lipid and carbohydrate. One way in which these diets differ is in their lipid composition(*42, 43*).

The yeast *Saccharomyces cerevisiae* contains predominantly palmitoleic acid (16:1) and oleic acid (18:1) and (like animals) does not appear to introduce unsaturated bonds beyond the delta 9 position(*42, 44*). In contrast, plants can introduce unsaturated bonds at the delta 12 and delta 15 positions, producing omega-3 and omega-6 fatty acids. Overall, the lipids present in plant food are longer and more unsaturated than those in yeast food. Also, plant food contains phytosterols, while yeast food contains fungal sterols(*42*).

Strikingly, we found that the different lipids present in plant and yeast food exert profound effects on membrane lipid composition, developmental rate, fertility and lifespan(*43*). Animals that feed on plant food incorporate longer and more unsaturated fatty acids into their membrane lipids than those fed on yeast food, and their membranes contain phytosterols rather than fungal sterols(*42*). They develop more slowly than animals fed with yeast, and the adults that emerge live longer. These differences in developmental rate and lifespan on plant and yeast foods are due to alterations in the levels of systemic insulin/IGF signaling (IIS). A yeast-derived lipid or lipid-soluble factor causes lipoproteins to accumulate on specific neurons in the brain that connect to Insulin Producing Cells (IPCs). When this happens, these neurons active IPC’s, causing them to release *Drosophila* insulin-like peptides (Dilps). Dilps elevate systemic IIS, speed development, increase fertility and shorten lifespan(*43, 45, 46*).

Why should yeast-derived lipid cues drive higher levels of Insulin signalling independent of the caloric content of the diet? We speculated that such a mechanism might have evolved to allow animals to maximally exploit summer blooms of fungi in the wild. Furthermore, we wondered whether the IIS-dependent increase in developmental rate might allow flies to thrive at higher temperatures. Conversely, since *Drosophila melanogaster* do not appear to be able to produce lipids with highly unsaturated fatty acids unless they consume plants, we wondered whether feeding on plant lipids might be important for flies to adapt their membrane biophysical properties to survive low temperatures.

To begin to investigate these questions, we asked whether *Drosophila* might alter their strong preference for yeast as a food source and substrate for egg laying(36–40) at low temperatures. Indeed, when females of the wild type strain OregonR are shifted to 15°C, they begin to lay eggs near plant food rather than yeast food (Figure 1A,B and S1 D,E), although their feeding preference is still for yeast. At 12°C, they also shift their feeding preference to plant food after a short hiatus (Figure 1C–E and S2).

**Figure 1:**
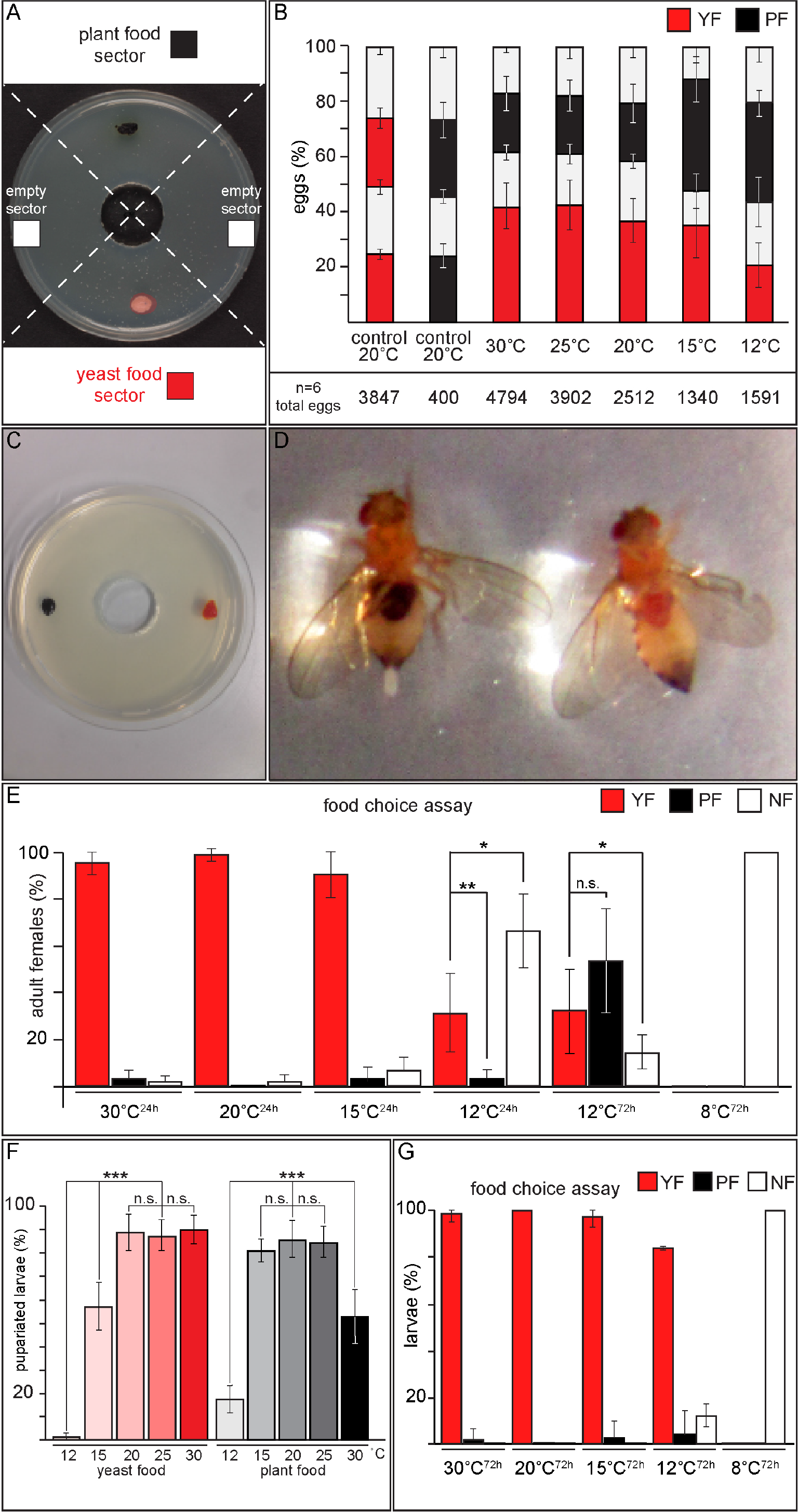
Food preference is temperature dependent. A. Females were allowed to lay eggs on plates divided into four sectors. Two sectors contained patches of either yeast or plant food and two intervening sectors were left empty B. Shows % eggs laid in each of the 4 plate sectors at the indicated temperatures, along with the total number of eggs counted from 6 plates representing 6 independent experiments. Within each bar, red indicates the plate sectors containing yeast, black indicates plate sectors containing plant food, and white indicates plate sectors with no food. As controls, we quantified eggs laid on plates containing identical foods on both sides. Error bars show standard deviation. C. Food-choice-plate with colored yeast (red) and plant food (dark blue). D. Color of food ingested is visible in abdomens of females that chose yeast (red) or plant food (dark blue). E.Percentages of females that had ingested yeast food (red), plant food (black), or no food (white) at the indicated temperatures after the indicated times (in hours). * indicates significant difference P values (Students t-test) are indicated: * = p<0.** = p<0.01, n.s. = not significant; Error bars show standard deviation. F.Percentages of larvae that successfully pupariate at specified temperatures when fed with yeast food (red scaled) or plant food (grey scaled). P values (Students t-test) are indicated: *** = p<0.001, n.s. = not significant; Error bars show standard deviation. G.Larval food preference was assayed using colored food (as in D). Plot shows the percentages of larvae choosing yeast food (red), plant food (black) or no food (white) at the indicated temperatures and time intervals. Error bars show standard deviation.

Do these different foods influence larval survival at different temperatures? To investigate this, we raised OregonR larvae on yeast or plant food at different temperatures and quantified the fraction of larvae reaching pupariation. Although 90% of yeast-fed larvae pupariate at temperatures between 20 and 30°C, only half pupariate at 15°C and none at 12°C. In contrast, plant-fed larvae efficiently pupariate between 15 and 25°C, and a substantial number of puparia form even at 12°C. However plant-fed larvae are much less successful at 30°C (Figure 1F). Thus, feeding with yeast improves high temperature development, whereas feeding with plants improves low temperature development. Larvae (unlike adults) do not change their feeding preference at low temperature - given a choice between plant and yeast food at 12-15°C, most larvae eat yeast and die (Figure 1G). This suggests that female preference for laying eggs near plant food at low temperature may be important for survival of their progeny below 15°C.

We next examined whether feeding with yeast or plant food influenced adult survival of OregonR at different temperatures. We first asked whether diet exerts similar effects on lifespan throughout the viable temperature range. In general, feeding on plants, rather than yeast, should increase *Drosophila* lifespan because IIS activity is lower in plant-fed animals - we previously showed that plant food extends lifespan when flies are kept at 25°C(*43, 46*). Temperature itself also influences lifespan(*47*). Like other poikilotherms, *Drosophila* live longer at cooler temperatures. Lifespan increases down to about 10°C, but lower temperatures are damaging and increase mortality(*48*). To ask how diet influences lifespan at different temperatures, we placed adult females that had been raised on normal food at 21-23°C in vials containing either yeast or plant food at 8, 12, 20 or 30°C, and monitored survival over time. Plant food extends lifespan at 8, 12 and 20°C. Maximal lifespan was observed at 12°C - at this temperature, plant-fed flies can live up to 6 months (Figure 2C). At 8°C the lifespan of both plant and yeast-fed flies decreases, but plant-feeding still increases lifespan (Figure 2D). Interestingly, there was no significant difference in lifespan of yeast and plant-fed flies at 30°C (Figure 2). Thus, reducing the activity of the insulin signalling pathway does not extend lifespan at high temperature.

**Figure 2.**
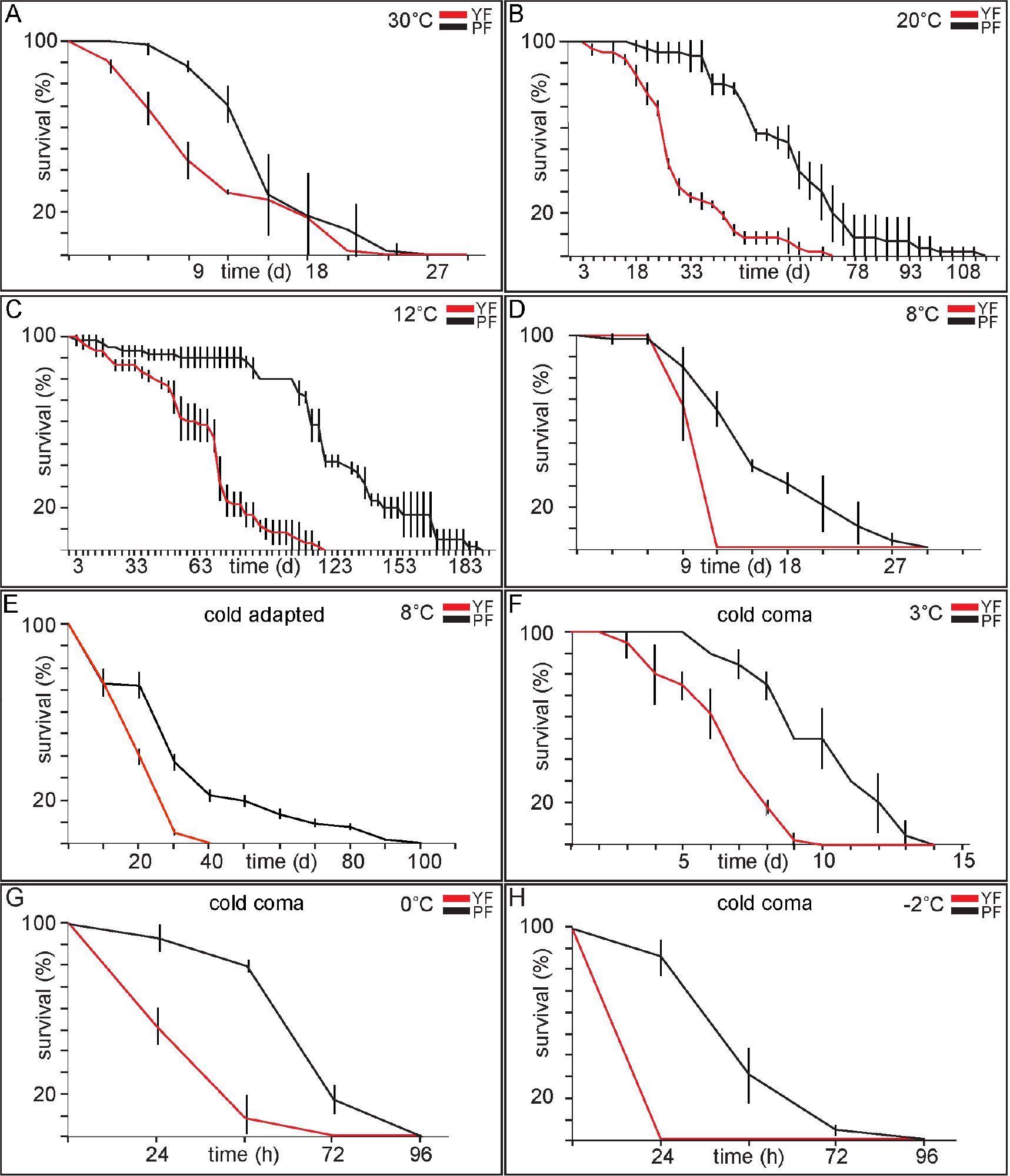
Plant food extends adult survival at low temperature. (A-H). Flies raised at 21-23°C(A-D,F-H) or at 15°C (E) on food containing both yeast and plant material were transferred after eclosion to vials containing either yeast (red) or plant (black) food for 10 days, then transferred to 15°C on the same food for two days. Flies were then transferred to the indicated temperatures. Panels show % females surviving at indicated times in days (A-F) or hours (G,H). Error bars show standard deviation.

While reduced insulin signalling may largely account for the longer lifespan of plant-fed flies at lower temperatures, we noticed that plant food also improved motor coordination between 12 and 8°C. Many of the yeast-fed flies we counted as alive in lifespan assays nevertheless appeared uncoordinated. Previous studies have shown that lack of coordination presages the onset of chill coma(*49*). To quantify coordination differences, we raised flies on plant or yeast food at 21-23°C, then placed them at either 15, 12 or 8°C for 72 hours. We then measured the time they required to climb up and out of a plastic tube at the same temperatures. This analysis confirmed that yeast-fed flies start to become uncoordinated at 12°C and are almost completely immobile at 8°C, while plant-fed animals remain mobile and capable of geotaxis at both temperatures (Figure 3A,B and supplementary movies A-H). Thus, plant food may also further improve survival at low temperature by maintaining motor function.

**Figure 3.**
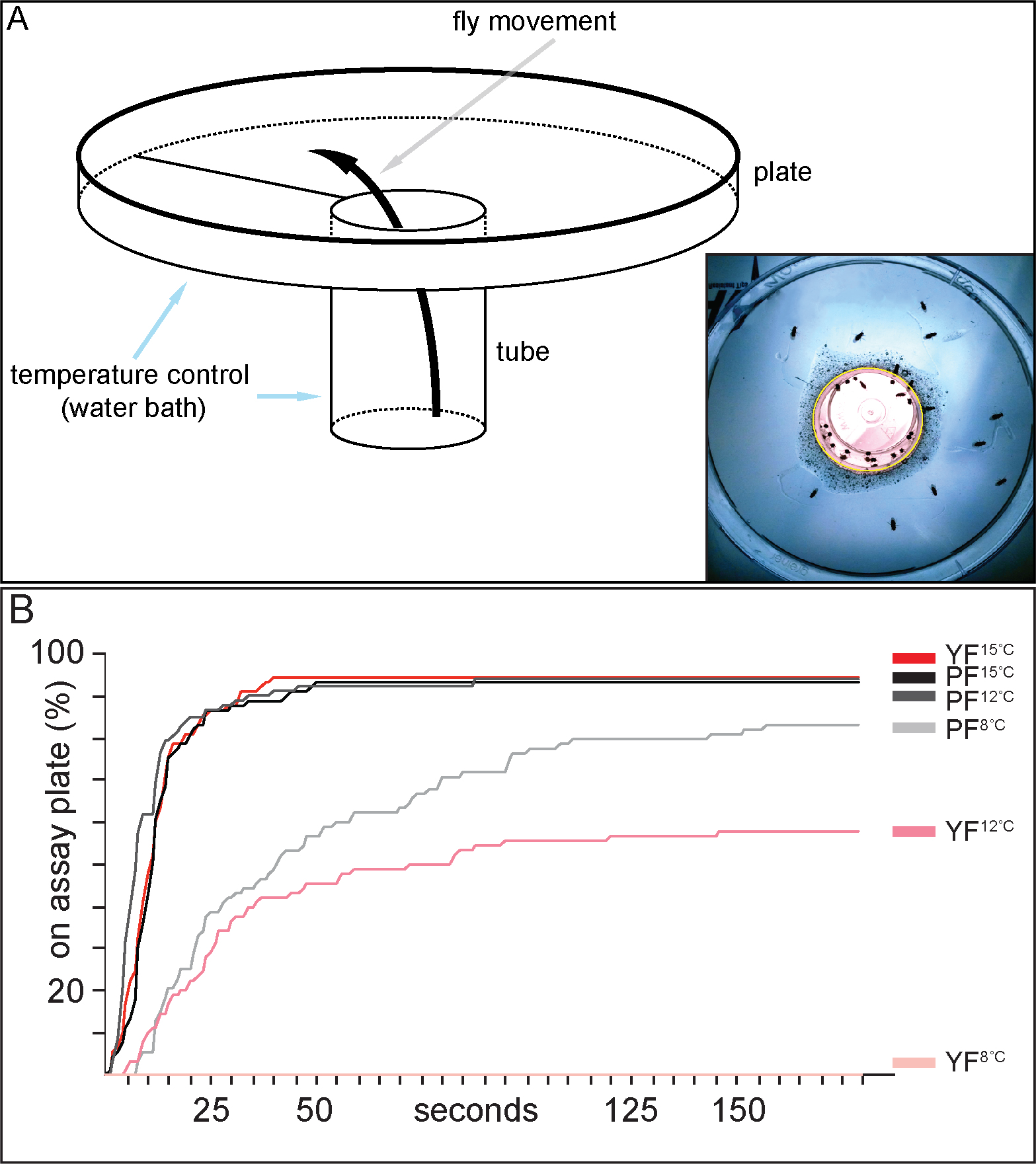
Temperature dependent mobility of adult flies. A. Cartoon of mobility assay plate and an inset showing a photograph of flies in such a plate. B. Flies were raised at 21-23°C on either yeast or plant food and than kept for 72h at 15, 12 or 8°C. These flies were placed on motility assay plates (shown in A) in a water bath at the corresponding temperatures, and filmed over the next 3 minutes. The length of time required for flies to crawl out of the tube and reach the plate was quantified in triplicate for each food and temperature condition. Red-scale indicates yeast food, greyscale indicates plant food.

At about 3°C and below, flies enter cold coma. When subjected to temperatures below −2°C, survival is typically a matter of hours(*17*). To investigate whether diet influenced survival at these temperatures, we subjected OregonR flies that had been raised on plant or yeast food at 21-23°C to cold coma at different temperatures for different lengths of time. We then quantified the number of flies that were able to awaken and survive for at least 24 hours afterwards (Figure 2F-H). Plant feeding almost doubles the median survival time between 3°C and −2°C.

Several studies have demonstrated that periods of pre-adaption to cold temperatures, or development at cooler temperatures, can increase cold tolerance of *Drosophila melanogaster(18, 50, 51)*. To ask whether a plant diet could further improve cold tolerance induced by previous exposure to cooler temperatures, we raised larvae on yeast or plant food at 15°C rather than 21°C and assayed survival of emerging adults at 8°C. Raising larvae at 15°C extends adult survival at 8°C independent of diet, but adults that fed as larvae on plant food live much longer than those fed with yeast (Figure 2E). Thus, the combination of a plant-based diet with pre-adaption to cooler temperatures increases cold tolerance more than either alone.

Thus far, we have investigated how *Drosophila* cope with temperature changes when temperature is held constant and flies experience a 12 hour: 12 hour light: dark cycle. However in nature, temperature fluctuates daily and day length decreases as winter approaches. We wondered how diet might influence development and cold tolerance under these more natural conditions. We therefore compared how plant food and yeast food influenced pupariation timing and adult survival of animals placed on the MPI-CBG rooftop in Dresden, Germany, between September 18^th^ 2015 and January 12^th^ 2016. During this time, diurnal temperature fluctuations ranged from 23°C/12°C to −1°C/−9°C (Figure 4A).

**Figure 4.**
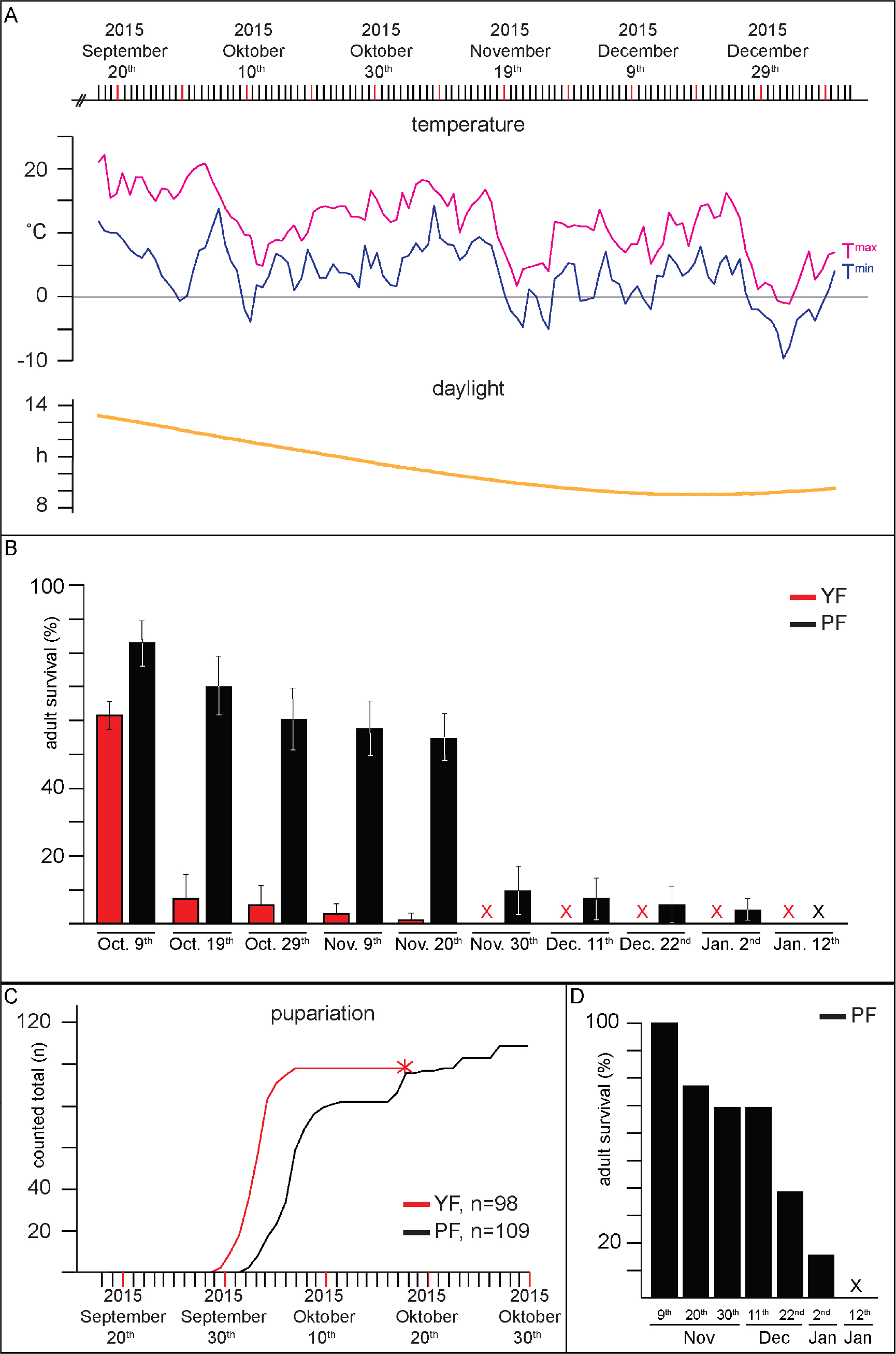
Plant food enables *Drosophila* to survive outdoor temperature fluctuations in the fall and winter. A. Upper panels show temperature curves (T^max^ magenta, T^min^ blue) and daylight time (middle, dark yellow) for Dresden, Germany (51.03N, 13.78E, 119m above sea level) from September 18^th^, 2015 through January 12^th^, 2016. B. Flies were raised on yeast or plant food at 21-23°C. 20 females and 15 males (aged between 14 and 21 days) transferred into 9 fresh food-tubes and placed outdoors (roof of the MPI-CBG) on September 18^th^. After collecting eggs for 3 days (see C), flies were inspected for viability and transferred to new food vials every 10 days. Shows the percent of flies surviving on the rooftop when fed with plant food (black) and yeast food (red), as of different dates. Error bars indicate standard deviation. C. Eggs laid by yeast or plant-fed flies (kept outdoors) were collected from September 18^th^-21^st^, and then transferred to fresh food vials. Visual inspection of these vials confirmed our previous observation that yeast-fed females normally lay 10-fold more eggs than plant-fed females (see figure S1A-C). However, practical considerations prevented us from counting the number of eggs laid by flies on the roof. Larval development and the number of pupae formed in each vial was tracked outdoors every two days until November 1^st^ (lower panel). The plot shows the number of pupae formed by yeast-fed (red) or plant-fed (black) larvae over time. D. Panel shows the survival of flies from the population that developed on plant food outside and eclosed between October 20^th^ and November 6^th^.

On September 18^th^, we allowed 20 female and 15 male OregonR flies per vial to lay eggs on plant food or on yeast food for three days, then monitored the number of pupae formed in each vial over time at intervals of 10 days. Although flies lay five-fold more eggs on yeast food than on plant food (Figure S1A-C), larvae in yeast vials gave rise to only 1/2 as many pupae. Thus, plant-fed larvae raised outdoors developed to pupae about 10-fold more successfully than yeast-fed larvae. Pupariation of yeast-fed larvae ceased shortly after the night-time temperature dropped to −4°C on October 12^th^, and none of these animals ever emerged as adults. In contrast, larvae growing in plant food vials continued to pupariate until October 29th, and 16 adult flies emerged from these vials at the beginning of November. Thus, only larvae that feed on plant food are able to complete development under seasonal conditions characteristic of Dresden in the fall (Figure 4C). Interestingly the flies that emerged in November showed increased melani-sation of the thorax (Figure S3), consistent with previous observations(*52, 53*). Increased melanisation in cold-blooded animals is thought to increase heat absorption and improve performance in the cold(*54, 55*).

Examining the developmental rates of plant and yeast-fed larvae raised outdoors revealed striking differences compared to developmental rates at constant temperature with the same average. The average of the fluctuating outdoor temperatures during the time that larvae were developing was 12.1 +/−1.6 °C. Interestingly plant-fed larvae raised outdoors began to pupariate after 10 days - about five times as fast as at a constant temperature of 12°C, and even three times as fast as at a constant temperature of 15°C. Thus, it would appear that plant-fed animals develop faster when exposed to diurnal temperature fluctuations outside than when exposed to the same, or even higher, constant average temperature. Yeast-fed animals also benefit from developing outdoors, though to a lesser degree. While yeast-fed larvae hardly pupariate at a constant temperature of 12°C, a fraction of these animals do pupariate when temperatures fluctuate around 12°C, and they develop about twice as fast as at a constant temperature of 15°C (Figure 4C). Previous studies have observed only small differences between larval development rates at constant versus fluctuating temperatures(*56*), although fluctuating temperature clearly increases developmental rates in other insects(*22*). It may be that other features of an outdoor environment, such as the irregularity of temperature fluctuations or decreasing day length, could speed *Drosophila* larval development, especially on plant food.

We next examined how diet influenced the survival of adults exposed to outdoor temperature changes. OregonR flies, aged between 10-22 days that had been raised on plant or yeast food at 21-23°C were placed in vials on the rooftop on September 18^th^, and their survival was monitored every 10 days (Figure 4B). Most yeast fed-animals died in the period between October 9^th^ and October 19^th^, when the temperature fell to −3°C on two consecutive nights. In contrast, 70% of plant-fed animals survived this temperature drop. Most survived until the interval between November 19^th^-29^th^, when temperatures fluctuated between 5 and −5°C for 5 days. Nevertheless, about 10% of plant-fed adults survived even this extended period of cold (Figure 4A,B). Interestingly, the flies that had developed on plant food outdoors and emerged during November survived the cold snap between November 19^th^ and 29^th^ much better that flies that had been raised inside at 21-23°C (Figure 4A,D). These animals, and those remaining from the first outdoor generation, died between January 2^nd^ and 12^th^, when day and night time temperatures dropped to −1/−9°C for three days (Figure 4A,B,D).

Taken together, these data show that feeding on plant food, but not yeast food, allows OregonR to survive outdoor conditions in Dresden up to midwinter. We note that *Drosophila melanogaster* are thought to seek sheltered over-wintering sites in the wild(*57, 58*) - an option not available to our experimental flies.

By what mechanisms do plant and yeast-based diets alter the viable temperature range? We first addressed how yeast food improved the success of larval development at high temperature. In general, higher temperature elevates metabolism and forces larvae to develop faster up until temperature approaches the edge of the viable range(*13*). Yeast-fed animals have much higher levels of IIS than plant-fed animals and develop at a faster rate(*43*). We therefore wondered whether higher levels of insulin signalling might extend the range over which the developmental rate increases with temperature. Indeed, yeast-fed larvae (which have higher IIS) develop slightly faster at 30°C than they do at 25°C, while plant-fed larvae develop more slowly at 30°C (Figure 5A). To investigate whether IIS was important for high temperature survival, we manipulated IPC activity and Dilp2 production in plant and yeast-fed larvae. Increasing Dilp2 production speeds development of plant-fed larvae and rescues their ability to pupariate at 30°C without changing pupariation efficiency at 15°C (Figure 5B). In contrast, further increasing Dilp2 production in yeast-fed larvae does not increase the speed of larval development at any temperature (Figure 5A), suggesting that IIS is already maximal. Consistent with this, over-expressing Dilp2 does not improve the already high success of pupariation of yeast-fed larvae at 30°C (Figure 5C). Strikingly, blocking IPC activity abrogates survival of both plant and yeast-fed larvae at 30°C, but still allows survival at 15°C (Figure 5C). Thus, high systemic IIS is necessary and sufficient for larval development at 30°C. However the converse is not true - low IIS does not improve and is not required for larval survival at low temperature.

**Figure 5.**
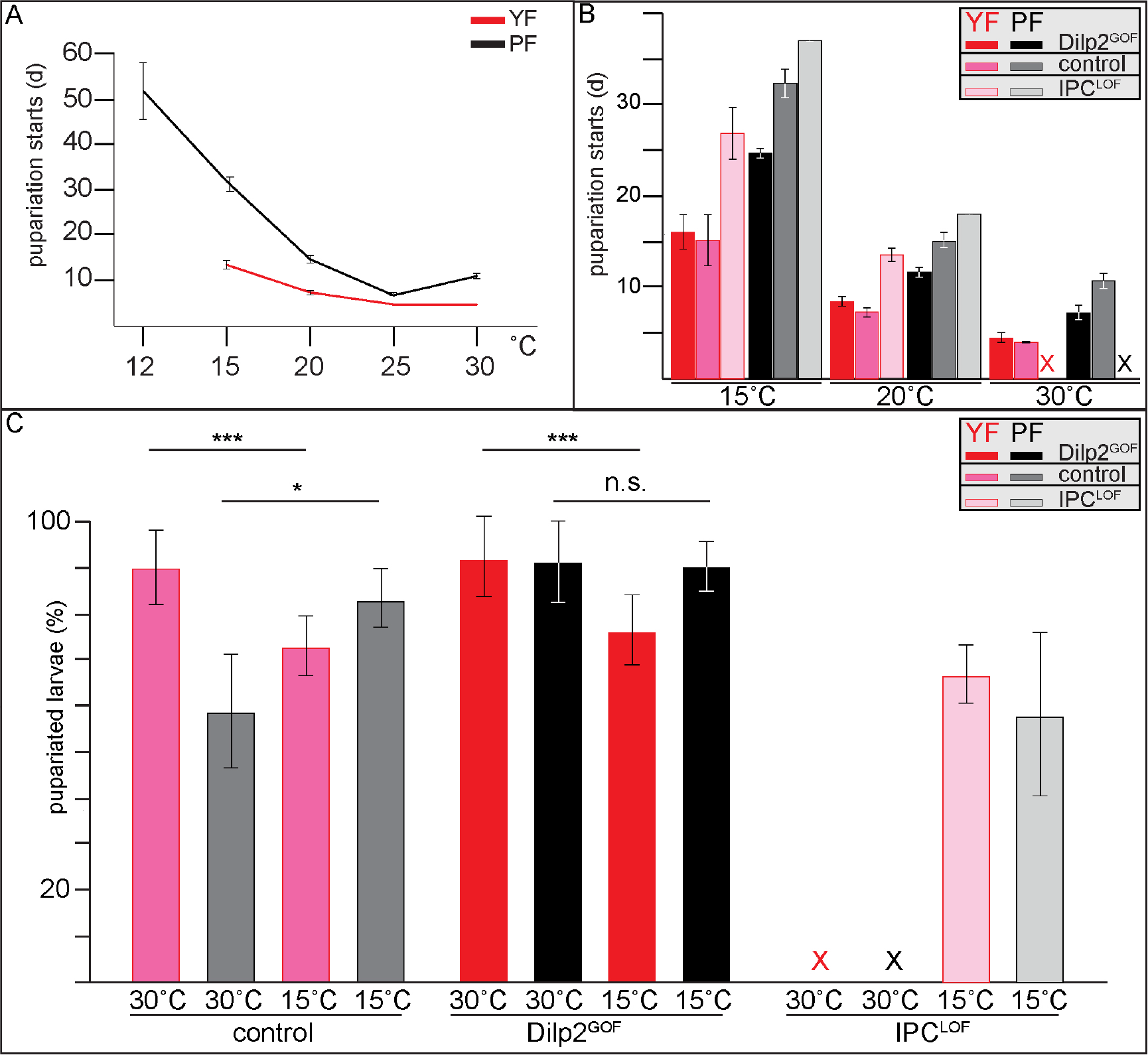
Insulin signaling is essential for survival at high temperatures. A. Shows the average time required for larvae to begin pupariating when fed with plant (black) or yeast (red) food at the indicated temperatures. Times are the averages from 3-5 biological replicates. Bars indicate standard deviations B. Panel shows the time required for larvae of indicated genotypes to begin pupariating on yeast food (red scale) and plant food (grey scale) at the indicated temperatures. We compared control larvae (*dilp2*Gal4/+) to larvae that over-express Dilp2 under the control of dilp2Gal4 (*Dilp2*^GOF^: *dilp2Gal4*>Dilp2), and to larvae in which IPC activity and Dilp secretion is inhibited by expression of a dominant negative potassium channel (IP-C^LOF^: *dilp2*Gal4>Kir2.1GFP). C. Panel shows the percent of larvae of the indicated genotypes that pupariate successfully on yeast food (red scale) and plant food (grey scale) at the indicated temperatures. Control: *dilp2*Gal4/+, Dilp2^GOF^: *dilp2*Gal4>Dilp2 and IPC^LOF^: *dilp2*Gal4>Kir2.1GFP. P-values (Students t-test) are indicated: *=p<0.025, *** = p<0.001, n.s. = not significant; Error bars show standard deviation.

What features of plant food promote low temperature survival? *Drosophila* require plant-derived unsaturated fatty acids to produce highly unsaturated membrane phospholipids(*42*). We therefore wondered whether plant feeding might be necessary for *Drosophila* to adapt its biophysical membrane properties to low temperature, consistent with models of homeoviscous adaption (*3*). To pursue this possibility, we first asked whether the lipid fraction of plant food was responsible for larval cold resistance. To do so, we devised diets containing the lipid-depleted water-soluble fraction of yeast, in combination with the lipid fraction derived from either yeast or plant food. We then raised larvae on these diets at different temperatures, and monitored pupariation efficiency (Figure 6A). Neither of the new foods were as effective as yeast food or plant food at supporting pupariation at any temperature - even at 20°C, only 40-60% of larvae pupariated (compared to 80-90% on full media). Thus, some nutrients may have been destroyed by the chloroform extraction required to produce lipid-depleted food and lipid extracts. Nevertheless, plant and yeast lipids still had strikingly different capacities to support larval development at different temperatures. At 30°C, larvae pupariated 8 times as often when fed with yeast lipids compared to plant lipids. This is consistent with our previous observation that a lipid extract from yeast is sufficient to active IPC neurons and elevate insulin signalling(*43*). At 15°C, larvae pupariated 7 times as often when fed plant lipids. Thus, plant lipids enhance low temperature development even when all other dietary components are derived from yeast, indicating that plant lipids are a key factor in the low temperature resistance of plant-fed animals.

**Figure 6.**
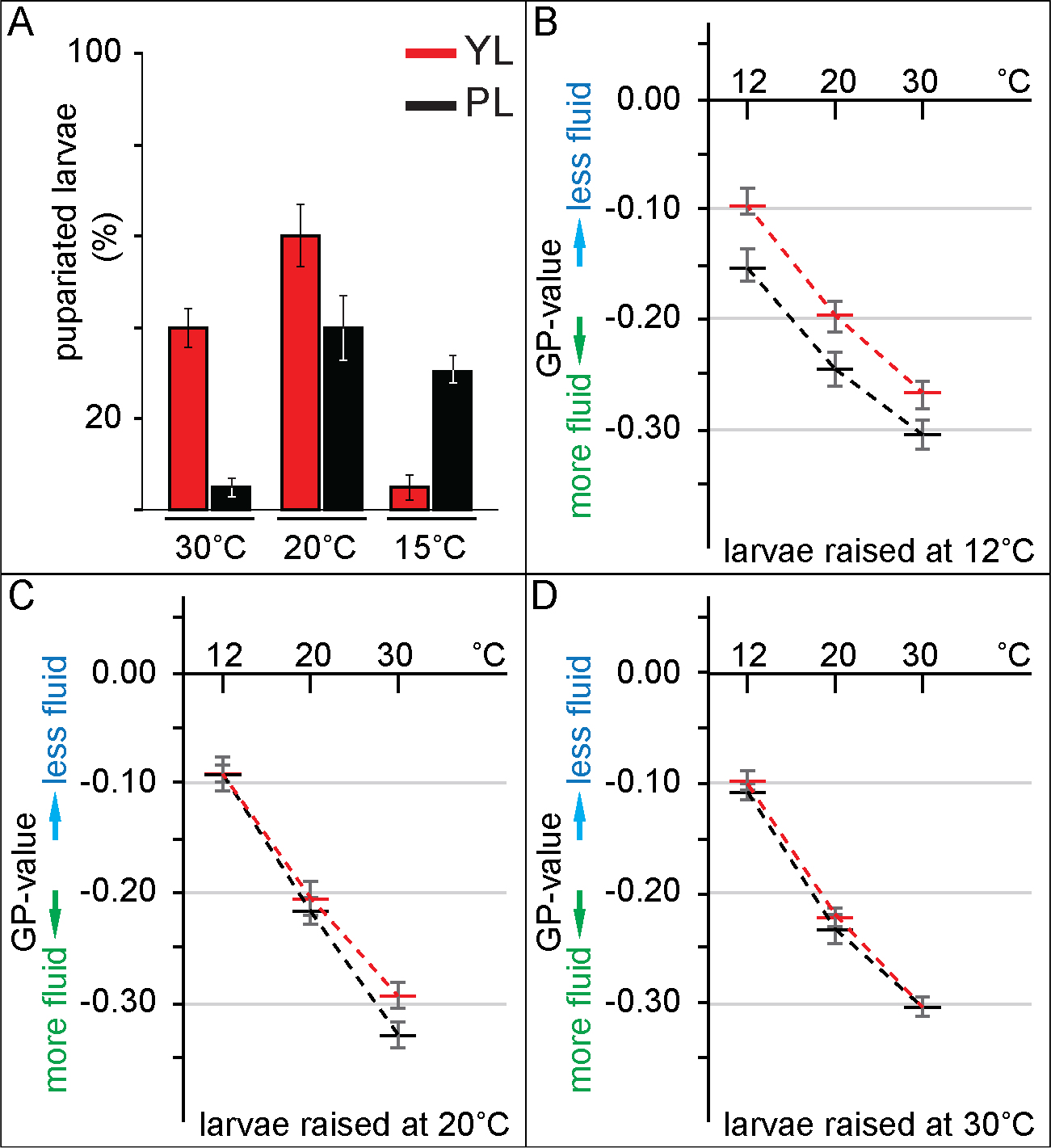
Dietary plant lipids decrease membrane order at low temperature. A. Panel shows the percentage of larvae pupariating successfully at the indicated temperatures when fed lipid-depleted food supplemented with lipid extracts of yeast (red) or plant (black) food at indicated temperatures. Error bars show standard deviation. B-D. Panels show average Generalized Polarization (GP)-values for Laurdan emission at 12, 20 or 30°C from liposomes prepared from lipid extracts of larvae raised on yeast food (red) or plant food (black) at 12°C (B), 20°C (C) or 30°C (C). Averages are from 3
biological replicates. Error bars (grey) show standard deviation. Lower GP values correspond to higher fluidity of membrane liposomes.

To investigate whether biophysical properties of membrane lipids varied with diet or with the temperature of development(*59*), we prepared liposomes from lipid extracts of plant and yeast-fed larvae raised between 12 and 30°C, and used C-Laurdan to measure their fluidity at different temperatures. C-Laurdan is a hydrophobic fluorescent dye that shifts its emission depending on the water content of lipid bilayers, a proxy for the degree of lipid order(*60, 61*). We found no significant differences in fluidity between liposomes prepared from plant and yeast-fed larvae raised at 20 and 30°C (Figure 6C,D). In striking contrast, membrane lipid fluidity of yeast and plant-fed animals raised at 12°C was very different (Figure 6B). While animals fed with yeast produced membrane lipids with similar fluidity at 12, 20 and 30°C, plant fed animals increased membrane fluidity when raised at 12°C. Three interesting conclusions can be drawn from these observations: 1) Despite their different lipidomes(*42*), plant and yeast-fed larvae raised between 20-30°C produce membranes with very similar biophysical properties. 2) *Drosophila* raised between 20 and 30°C do not appear to adapt the biophysical properties of their membrane lipids to temperature. Surprisingly, this suggests that flies living at 30°C have more fluid membranes than those living at 20°C. 3) Finally, *Drosophila* adapt membrane biophysical properties to reduce lipid order only if they feed on plants and only when raised at 12°C. Thus, the temperature at which plant-fed animals increase membrane fluidity coincides with the temperature at which plant-feeding is necessary to enhance larval survival (Figs. 1 and 6). This suggests that feeding on plants may promote survival at low temperature by allowing animals to increase fluidity of membrane lipids.

## Discussion

The mechanisms that allow *Drosophila melanogaster* to survive the winter in different climates are not completely understood. Here, we have shown that the common lab strain OregonR changes its feeding and egg laying preference from yeast to plant material at temperatures below 15°C, and that yeast and plant diets help animals survive heat and cold, respectively. OregonR was established over 90 years ago from flies caught in Roseburg Oregon, where the coldest average temperature in winter is 8°C. Since then, they have been maintained in many different laboratories where they are generally exposed to temperatures between 16 and 30°C. Despite the fact that this strain has not been forced to cope with extreme low temperatures for many years, feeding on plant food allowed OregonR to survive outside in Dresden from September through January, coping with several successive nights where temperature dropped to −5°C. Thus it seems likely that a dietary switch from yeast to plants would have allowed the flies from which OregonR was derived to survive the winter in their original environment even without seeking sheltered overwintering sites. It will be extremely interesting to investigate whether stocks that have been more recently established from colder climates exhibit the same switch in dietary preference, and whether a plant-based diet in the context of these different genetic backgrounds might allow even more striking cold resistance. Finally, it will be important to discover whether wild *Drosophila* feed more on plants when temperature drops, and if so, which plants they might favour. While dietary manipulations such as supplementation with cholesterol(*62*) or pro-line(*63*) have been observed to increase insect cold-resistance, a switch in feeding preference from yeast to plants is a mechanism that might plausibly function in the wild.

To what extent *Drosophila melanogaster* engages in homeoviscous adaption has been unclear. Although some changes in lipid composition have been observed at lower temperatures(*27–32*), membrane fluidity changes have not been measured in this organism. Our measurements of membrane fluidity show that *Drosophila melanogaster* can indeed modulate the biophysical properties of their membranes to increase fluidity when temperatures drop, but only if they are allowed to feed on plant material. Interestingly, this adaption does not occur throughout the viable temperature range. Lipids isolated from larvae raised at 20 and 30°C show no significant difference in their physical properties - liposomes prepared from these lipids increase fluidity with measurement temperature in exactly the same way. The fact that larvae do not appear to tune their lipid properties between 20°C and 30°C suggests that these animals simply allow membrane fluidity to increase with temperature in this range, although of course this remains to be tested *in vivo*. Since metabolism and development run faster at 30°C than at 20°C, it may be that proportional increases in membrane fluidity help keep biological processes coordinated at different temperatures. Altering membrane biophysical properties may only become important below a low temperature threshold where it is essential to prevent phase separation.

We were surprised to observe such similar biophysical properties of membrane lipids isolated from plant and yeast-fed animals between 20-30°C, because their membrane lipid composition is strikingly different(*42*). When larvae are raised at 25°C on plant or yeast food, membrane lipids in all tissues of plant-fed larvae contain more unsaturated fatty acids that those of yeast-fed larvae. In the brain, for example, the average number of unsaturated bonds per fatty acid increases by 30% on plant food. These differences in unsaturation are larger than those observed upon cold hardening in *Drosophila,* which are on the order of 1%(*64*). Despite such large differences in average fatty acid unsaturation, liposomes from plant and yeast-fed animals have very similar fluidity when prepared from larvae raised between 20 and 30°C. Thus, even large changes in fatty acid unsaturation do not necessarily lead to membrane fluidity differences. In addition to differences in fatty acid composition, plant and yeast-based diets also change the ratios of phospholipid head-groups, and alter the species and amounts of membrane sterols(*42*). Any of these changes could influence membrane biophysical properties. Thus it seems clear that the fluidity of biological membranes depends on combined features of many different lipids. The fact that plant and yeast fed animals can produce membranes with indistinguishable biophysical properties using very different populations of lipids suggests a remarkable ability to sense and tune these properties. Nevertheless, the unsaturated fatty acids provided by plants may allow *Drosophila* to tune membrane biophysical properties over a broader range.

It has been shown for many cold-blooded animals that the upper temperature limit for successful development occurs near the region where the developmental rate no longer increases with temperature. However, the causal relationship between developmental slowdown and lethality was difficult to prove, and what might limit the developmental rate at excessively high temperature was not known. We show that differences in the level of systemic insulin signalling set this limit and are responsible for the different developmental temperature limits of larvae fed on yeast and plant food. This finding supports the idea that the inability to continue to increase the developmental rate with temperature is a key limitation that sets the viable temperature range. How might Insulin signalling affect the relationship between growth and temperature? If increasing temperature drives biological reactions faster, it may be that the rate at which cells are supplied with the raw materials for growth and metabolism becomes limiting. Insulin and *Drosophila* Insulin-like peptides are potent anabolic hormones that promote cellular nutrient uptake, and the incorporation of sugars, lipids and amino acids into cellular constituents and storage forms(*65*). We would propose that a yeast diet, which promotes high levels of insulin signalling, allows nutrient uptake to keep pace with biological reactions driven faster by temperature.

Interestingly, distinct alleles of the *Drosophila* Insulin receptor have been shown to vary in frequency according to latitude and even according to season. These alleles have different activities and are associated with differences in developmental timing and body weight, as well as different adult sensitivities to cold and heat shock(*66*). It will be interesting to see whether these alleles also influence the temperature range over which development is successful.

Taken together, our observations support a testable model for *Drosophila* seasonal adaption. In summer, *Drosophila* feed on yeast, increasing systemic insulin signalling. This increases fertility and speeds development to allow exploitation of a transient resource, and also extends the upper temperature limit at which development can occur. In the fall, *Drosophila* begin to feed on and lay eggs near plant material. This increases the success of larval development at low temperature and prepares emerging adults for overwintering by preserving membrane fluidity and extending lifespan.

Seasonal variations in diet and fertility also occur in mammalian animal populations, and it has been known for some time that plant and animal fatty acids have different effects on mammalian insulin signalling(67–69). Our findings suggest that it may be productive to think about these differences in the context of seasonal adaption.

## Author contributions

Conceptualization, M.B., S.E., U.C.; Investigation, M.B., T.G., B.B. and M.G.; Writing - Original Draft, M.B. and S.E.; Writing - Review & Editing, U.C., M.B. and S.E.; Funding Acquisition, U.C. and S.E.; Resources, U.C. and S.E.; Supervision, U.C. and S.E.

## Acknowledgements

We thank Tony Hyman, Teymuras Kurzchalia, Kai Simons, Pavel Tomancak and Marino Zerial for critical comments on the manuscript. Ali Mahmoud provided expert technical assistance. This work was supported by funding from the Max Planck Society (SE and MB), the Deutsche Forschungsgemeinschaft Transregio 83, TP18 and TP19 (SE, MB, UC), Dresden International Graduate School for Biomedicine and Bioengineering DIGS-BB (TG) and from a German Federal Ministry of Education and Research grant to the German Center for Diabetes Research DZD e.V., (UC, MG, BB).

---

**Figure S1:**
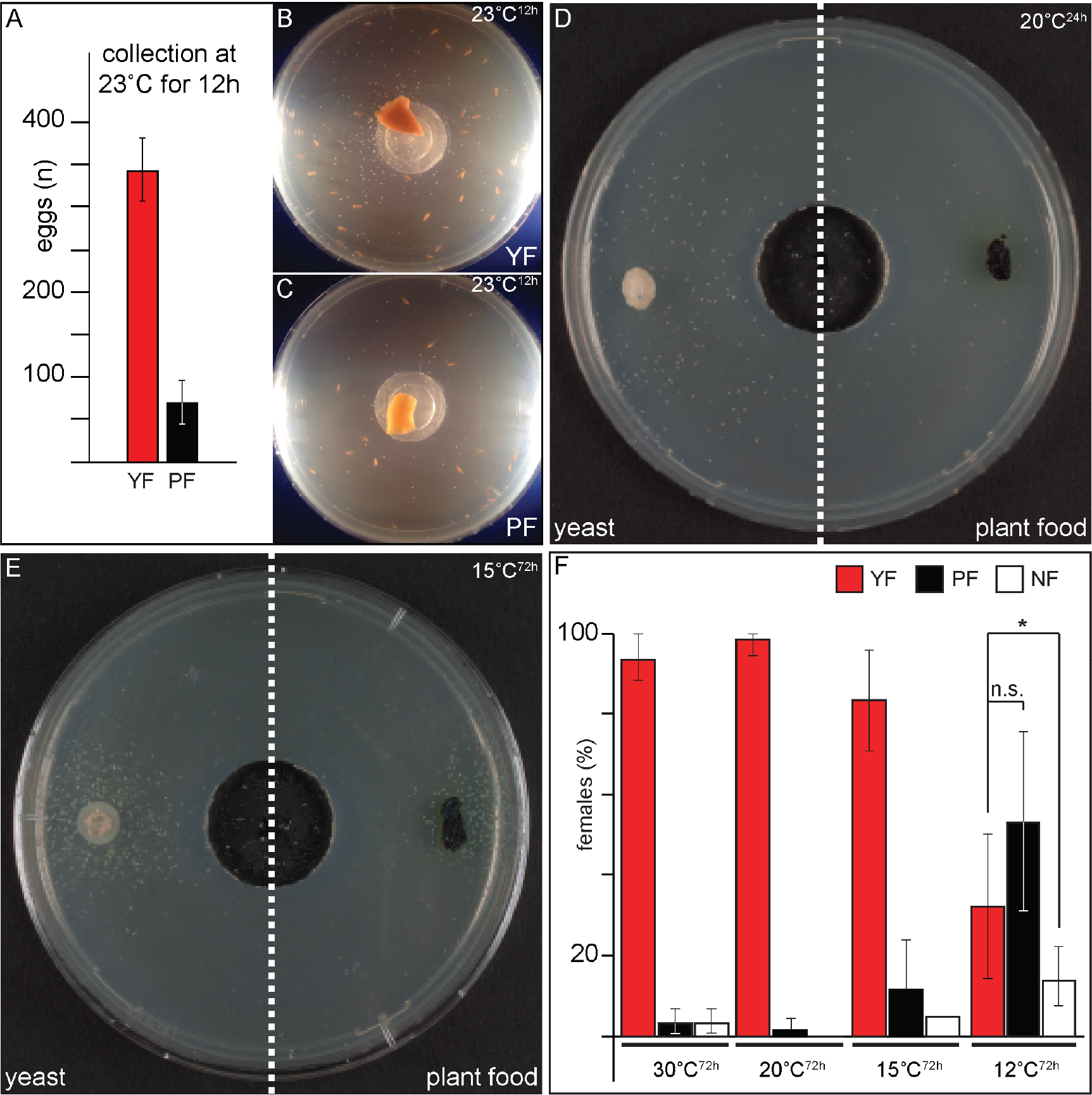
Food preference is temperature dependent. A. Numbers of eggs laid over a 12h period at 23°C by wild type flies (20 females, 15 males) kept on yeast food (red) or plant food (black) plates (B). Error bars indicate standard deviation. B, C Panels show representative photographs of egg collection plates with yeast (B) and plant food (C). D,E. show egg collection plates containing yeast food on the left and plant food on the right photographed after 24h at 20°C (D) or after 72h at 15°C (E). F. Female food preference was assayed at specified temperatures and indicated time intervals (X-axis). Plotted are flies (Y-axis, in %) that had consumed yeast food (red), plant food (black) or no food (white). Error bars show standard deviation.

**Figure S2:**
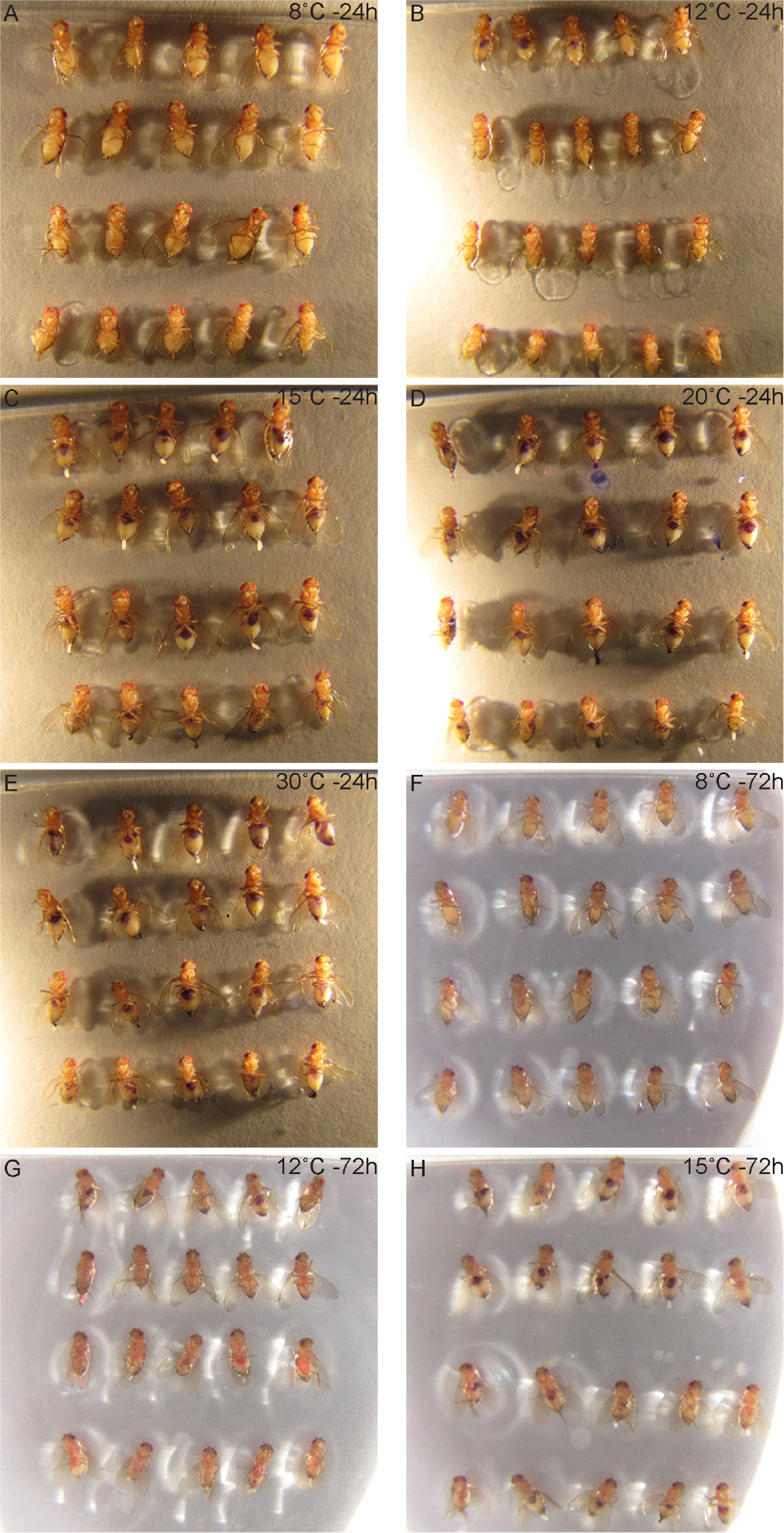
Mated females prefer plant food at 12°C. A-H. Photos show examples of raw data that are summarized in Figure 1D. Females given the choice between yeast food (blue) or plant food (red) at the specified temperatures were glued to a glass slide by the dorsal thorax (abdomen up) in arrays. Note that abdominal staining shows ingested food color.

**Figure S3:**
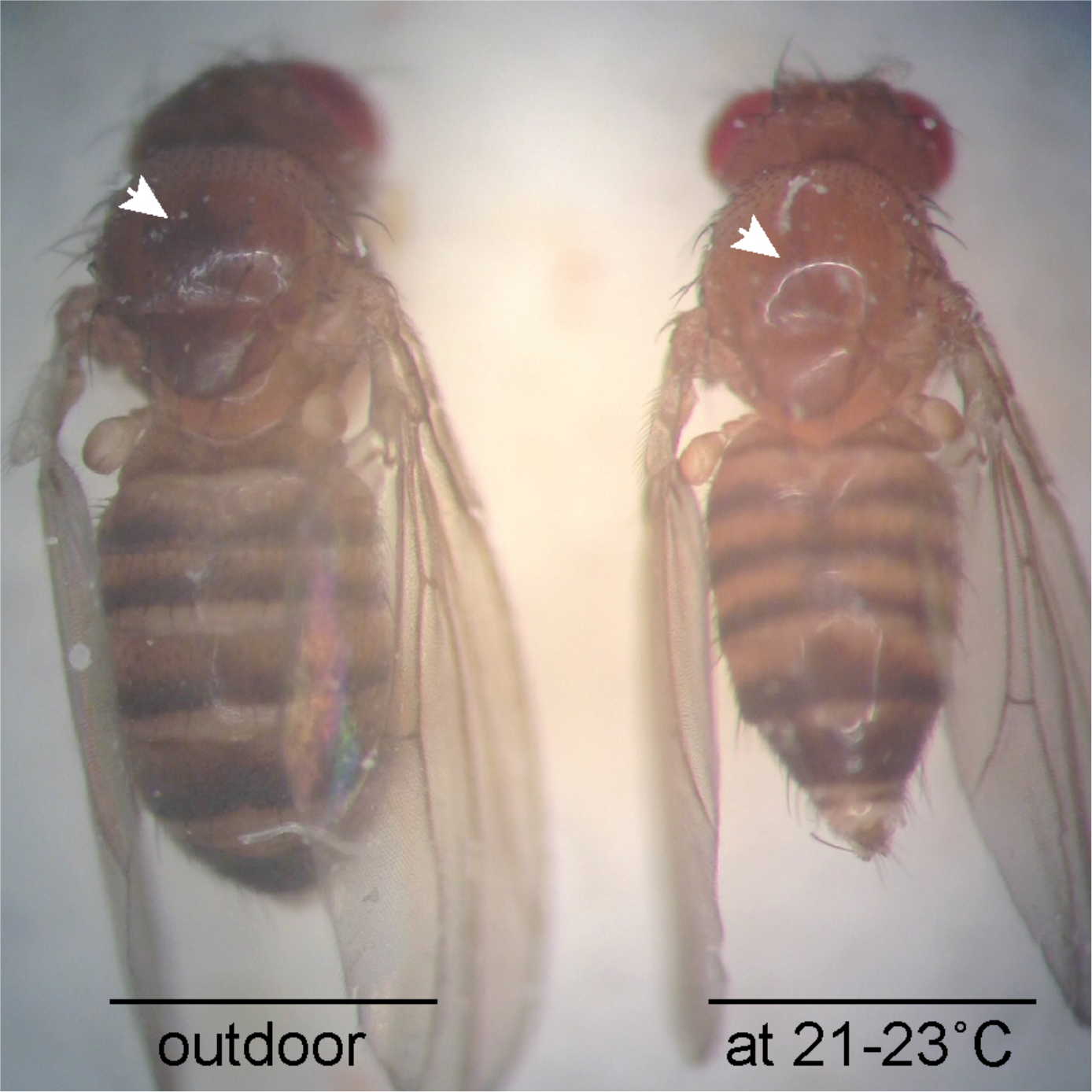
Fly pigmentation is temperature dependent. Photograph shows adult female flies raised on plant food either at 21-23°C (right side) or outdoors (from 18^th^ September - November 9^th^, left side). Arrow indicates increased melanisation on the thorax.

## Movie legends

A-F. show representative movies of mobility assays quantified in Figure 3B. (B,D,F) show plant-fed flies, (A,C,E) show yeast fed flies. Experiments were run at 15, 12 and 8°C as indicated.

## Experimental procedures

Fly stocks

If not indicated otherwise, flies were rised on YF under a 12h light/12h dark cycle at 21-23°C. oregonR (#5), dilp2Gal4 (#37516), UAS:Dilp2 (E. Hafen) and UAS:Kir2.1GFP (#6596) are available from Bloomington Stock Center.

### Behavioral assays

#### Egg laying assay

10d after eclosion 40 mated females and 20 males were transferred onto conditioned food choice assay plates and shifted to specified breeding conditions. Assay-plates were collected after 12h (30 and 20°C), 48h (15°C) and 72h (12°C), equatorially divided (each sector with respective food) and manually assayed for embryo numbers.

#### Food choice assay (adults)

Yeast-pellet (K classic yeast, Kaufland) suspended in 5ml 10% Glucose, incubated for 30’ at RT, stained with Coomasie blue G-250 (Sigma-Aldrich) and stored at 3°C (>24h). Plant food paste was stained with Ponceau S (Sigma-Aldrich) and stored at 3°C (>24h). Opposing each other, food types were placed in defined rectangular areas (5×10mm) 40mm distant from agar plate center (1% Agar, Greiner, circular 100×15mm).

Flies were raised at 21-23°C on normal food and hatched virgin animals transferred on YF. 10d after eclosion 20 mated females and 10 males were transferred onto conditioned assay plates, and kept on respective temperatures for 24h or 72h respectively. Collected flies were photographed and manually assayed for predominant gut color produced by food ingestion.

#### Food choice assay (larvae)

Embryos were collected in 1h intervals at 21-23°C, aged for 24h and hatched 1^st^ instar larva transferred onto preconditioned assay plates (see food choice assay (adults)). Samples were manually assayed for predominant gut color after 72h (exception: larva kept at 12°C or 8°C were sampled after 216h).

#### Motility assay

Flies were raised on PF or YF at 21-23°C and 12d after eclosion transferred to specified temperatures for 72h. Thereafter, motility and coordination was video-recorded in temperature controlled assay chambers. In brief, the used motility assay chamber consists of a 3cm tube (d=1cm) connected to a petri-dish (d=6cm). Flies were transferred into the tube und shifted for 15min to 8°C. After vortex at max speed for 5 sec assay platform was transferred into recording system at specified temperature. In the assay plate flies are forced to climb up the tube and than are allowed to explore the plane plate area.

### Survival assays

#### Pupariation assay

1h embryo collections were aged for 24h at 21-23°C. Hatched 1^st^ instar larva (20/vial) were transferred onto respective food and kept under specified breeding conditions. Each vial was assayed for pupariation manually in 24h intervals.

#### Lifespan assay

Flies were rised on normal food at 21-23°C or 15°C (cold adapted flies) and eclosed animals transferred onto YF or PF. After 10d mated females (20) and males (10) were transferred onto respective fresh cooled (15°C) food vials and kept at this temperature for 48h. Subsequently vials were shifted to specified breeding conditions. Surviving females were counted in 72h intervals and transferred onto fresh food if required.

#### Lipid rescue assay

Wild type embryos were collected in 1h intervals at 21-23°C, transferred onto fresh apple-juice agar plates and after 24h hatched 1^st^ instar larvae were transferred onto lipid supplemented LD-food at specified temperature. Assayed were formed pupae.

### Biochemistry

#### Lipid extraction

For each lipid sample thirty 3^rd^ instar wild type larvae raised at specified conditions were dissected (on ice in HEPES-buffered saline (HBS): 25mM HEPES, 150mM NaCl, pH 7.25) to remove the gut and CNS, snap-frozen in liquid nitrogen and stored at −80°C until lipid extraction. Tissue samples were thawed on ice, homogenized in HBS using an IKA^®^ ULTRA-TURRAX^®^ disperser (level 5, 1min), and lipid-extracted by the BUME method (Lofgren, 2012 #450). Extracted lipids were stored in chloroform / methanol (2:1) solution at −80°C.

#### Liposome preparation

A volume containing 40 nmol phospholipids was dried under a stream of pressured air and left under vacuum for 2 h to ensure complete removal of solvents. The lipid film was rehydrated with 200¼l HBS with vigorous shaking for 30min, followed by 10 freeze-thaw cycles, and extruded through a 100nm filter to obtain homogeneous 100nm unilamellar liposomes.

#### Fluorescence spectroscopy

Membrane order was assessed by determining C-laurdan generalized polarization (GP) indices of liposomes prepared from fly lipids. Liposomes (200¼M phospholipids) were stained with C-laurdan (0.4¼M) and incubated for 30min in the dark at room temperature. Fluorescence spectra were obtained with a FluoroMax-4 spectrofluorometer (Horiba) equipped with a temperature-controlled Peltier element (Newport) at 12, 20, and 30°C. Excitation was 385nm and emission was recorded at 400-600nm with 1nm. Spectra were background-corrected and GP values were calculated as described in (Kaiser, 2009 #235).

